# Genetic regulators of mineral amount in Nelore cattle muscle predicted by a new co-expression and regulatory impact factor approach

**DOI:** 10.1101/804419

**Authors:** Juliana Afonso, Marina Rufino Salinas Fortes, Antonio Reverter, Wellison Jarles da Silva Diniz, Aline Silva Mello Cesar, Andressa Oliveira de Lima, Juliana Petrini, Marcela M. de Souza, Luiz Lehmann Coutinho, Gerson Barreto Mourão, Adhemar Zerlotini, Caio Fernando Gromboni, Ana Rita Araújo Nogueira, Luciana Correia de Almeida Regitano

## Abstract

Mineral amount in bovine muscle affect meat quality, growth, health and reproductive traits in beef cattle. To better understand the genetic basis of this phenotype, we implemented new applications of use for two complementary algorithms: the partial correlation and information theory (PCIT) and the regulatory impact factor (RIF), by including GEBVs as part of the input. We used PCIT to determine putative regulatory relationships based on significant associations between gene expression and mineral amount. Then, RIF was used to determine the regulatory impact of genes and miRNA expression over mineral amount. We also investigated over-represented pathways, as well as evidences from previous studies carried in the same population, to determine regulatory genes for mineral amount *e.g. NOX1*, whose expression was positively correlated to Zn and was described as regulated by this mineral in humans. With this methodology, we were able to identify genes, miRNAs and pathways not yet described as important for mineral amount. The results support the hypothesis that extracellular matrix interactions are the core regulator of mineral amount in muscle cells. Putative regulators described here add information to this hypothesis, expanding the molecular relationships between gene expression and minerals.

## Introduction

Mineral amount affects meat quality [1]–[4], reproduction [5], health and growth performance [6], [7] in beef cattle and the control of mineral homeostasis depends on genetic factors, among others [8]. Understanding the genetic aspects linked to mineral amount in bovine muscle can lead to a better modulation of this trait, allowing for future production of healthier, more productive animals, and better-quality meat.

A differential expression approach detects genes and pathways underlying mineral amount in Nelore cattle, by comparing extremes of the population used herein [9] [10]. However, as mineral amount traits occur in a continuous distribution, to verify these relationships and infer regulatory modes of action, it is necessary to study the whole population. It is possible to go beyond contrasting extreme phenotypes, beyond differential expression [11]. Thus, by applying a co-expression network approach it is possible to identify genome-wide genes with similar expression patterns related to specific phenotypes or conditions. In this methodology, traits are usually integrated into the analysis in a condition-dependent network, by previous selection of genes or sample clusters related to the trait before the analysis [12]. Another way of including phenotypes to select gene groups putatively involved with them, already used for mineral amount in our population [13], is to cluster all expressed genes by their co-expression profiles and then associate these clusters to the phenotypes using the Weighted correlation network analysis (WGCNA) iR package [14]. In this case, groups of genes with similar functions are identified and associated with the phenotypes.

Among the challenges of these methods regarding phenotype inclusion is that no single approach is used to search genome-wide for specific genes linked to phenotypes without prior selection. Also, it is challenging to pinpoint the direction of interactions or the regulation, as co-expression networks do not provide this information a priori [12]. To overcome these limitations, we propose a new application of the partial correlation and information theory (PCIT) algorithm, originally used for deriving gene co-expression networks, by identifying significant associations between expression profiles [15].

Additionally, we propose a new application of the regulatory impact factor (RIF) algorithm [16] to identify significant genes and miRNAs expression with regulatory impact over mineral amount in bovine muscle. To this end, we used the expression values of genes and miRNAs correlated to minerals instead of transcription factors (TFs) used in the original application, allowing the regulatory role to go beyond current functional annotation of the cattle genome. When calculating the RIF of genes and miRNAs with expression correlated with a mineral over the amount of this mineral, the mineral mass fraction genomic estimates of breeding values (GEBVs) were used instead of the expression data of selected gene.

Therefore, we were able to use GEBVs on the networks to identify regulatory elements linked to the phenotypes. This new use of the PCIT-RIF algorithms identified genes and miRNAs expression related to the mass fraction of calcium (Ca), copper (Cu), potassium (K), magnesium (Mg), sodium (Na), phosphorus (P), sulfur (S), selenium (Se), zinc (Zn) and iron (Fe) in Nelore steers’ *Longissimus thoracis* muscle. In short, we aimed to predict the regulatory impact of genes and miRNAs expression over mineral amount in Nelore muscle.

## Results

### Genes and miRNAs with expression values correlated to minerals

After data quality control, filtering, normalization and batch effect correction performed separately in the mRNA-Seq and miRNA-Seq from 113 samples, the expression of 12,943 genes and 705 miRNAs remained for further analyses. To identify genes and miRNAs with expression values correlated to ten different minerals, we carried out two different PCIT analyses, using our new application: i) PCIT general: incorporating genes, miRNAs expression and GEBVs together, and ii) PCIT miRNA: considering only miRNAs expression and GEBVs together. Simultaneously considering the results of both PCIT analyses, we identified a total of 242 genes and 35 miRNAs with expression values correlated to at least one mineral GEBV. From these, the expression of 46 genes and 12 miRNAs was correlated to more than one mineral GEBV. The number of genes and miRNAs with expression values correlated to each mineral ranged from 19 to 55 and from five to nine, respectively. The number of miRNAs’ expression that were correlated to a mineral in both PCIT analyses varies from zero to three (Table 1). There were two genes and one miRNA expression values correlated to six minerals, Vitamin D3 receptor (*VDR)* and bta-miR-92b correlated to Ca, K, Mg, Na, P and S; and Doublecortin (*DCX)*, correlated to K, Mg, Na, P, S, and Zn. From these analyses, we identified significant correlations among minerals’ GEBVs. There were no significant correlations between Se and other minerals (Figure 1). Correlations identified among K, Mg, Na, Zn, S, and P GEBVs ranged from 0.77 to 0.97.

**Figure 1.**
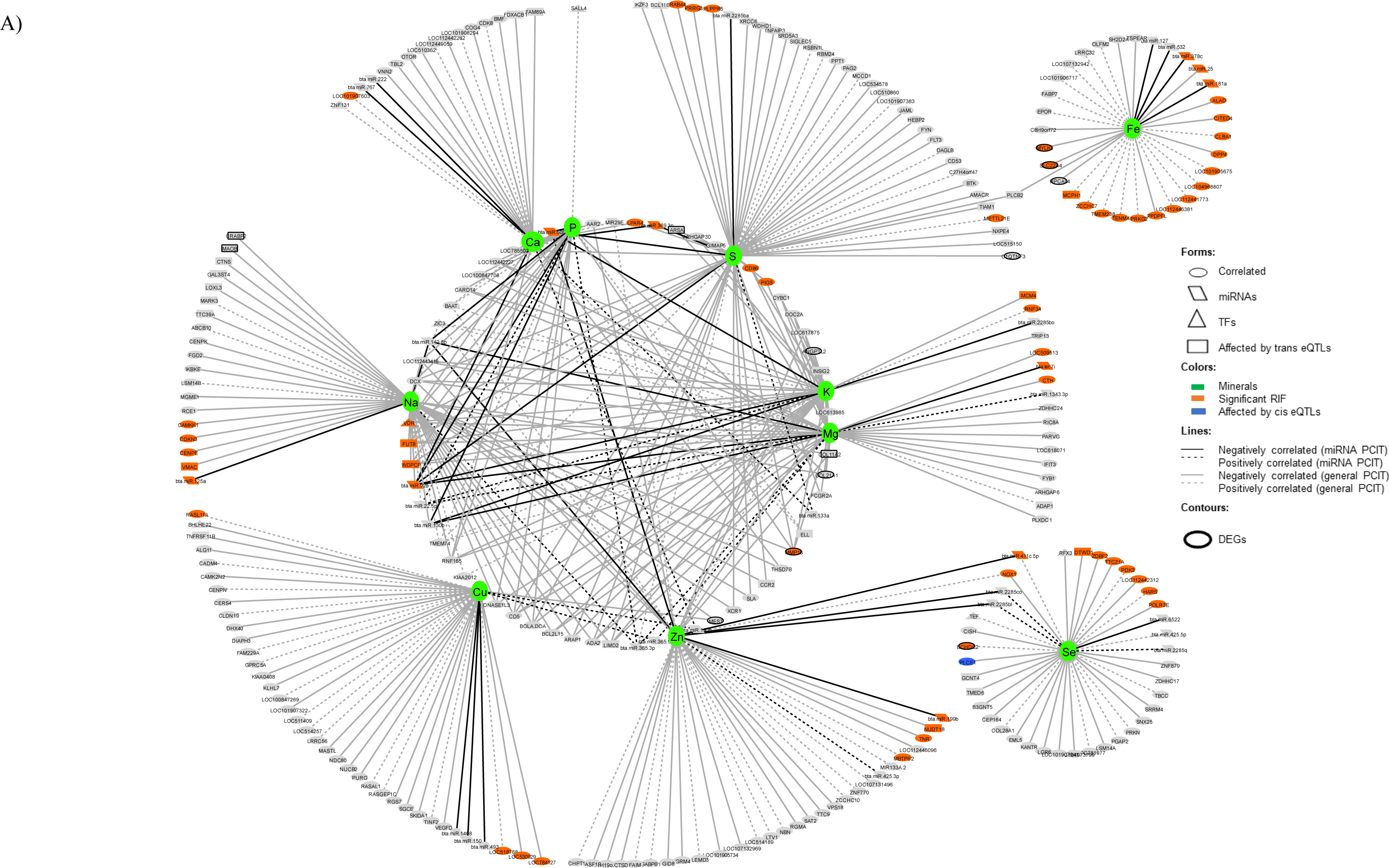
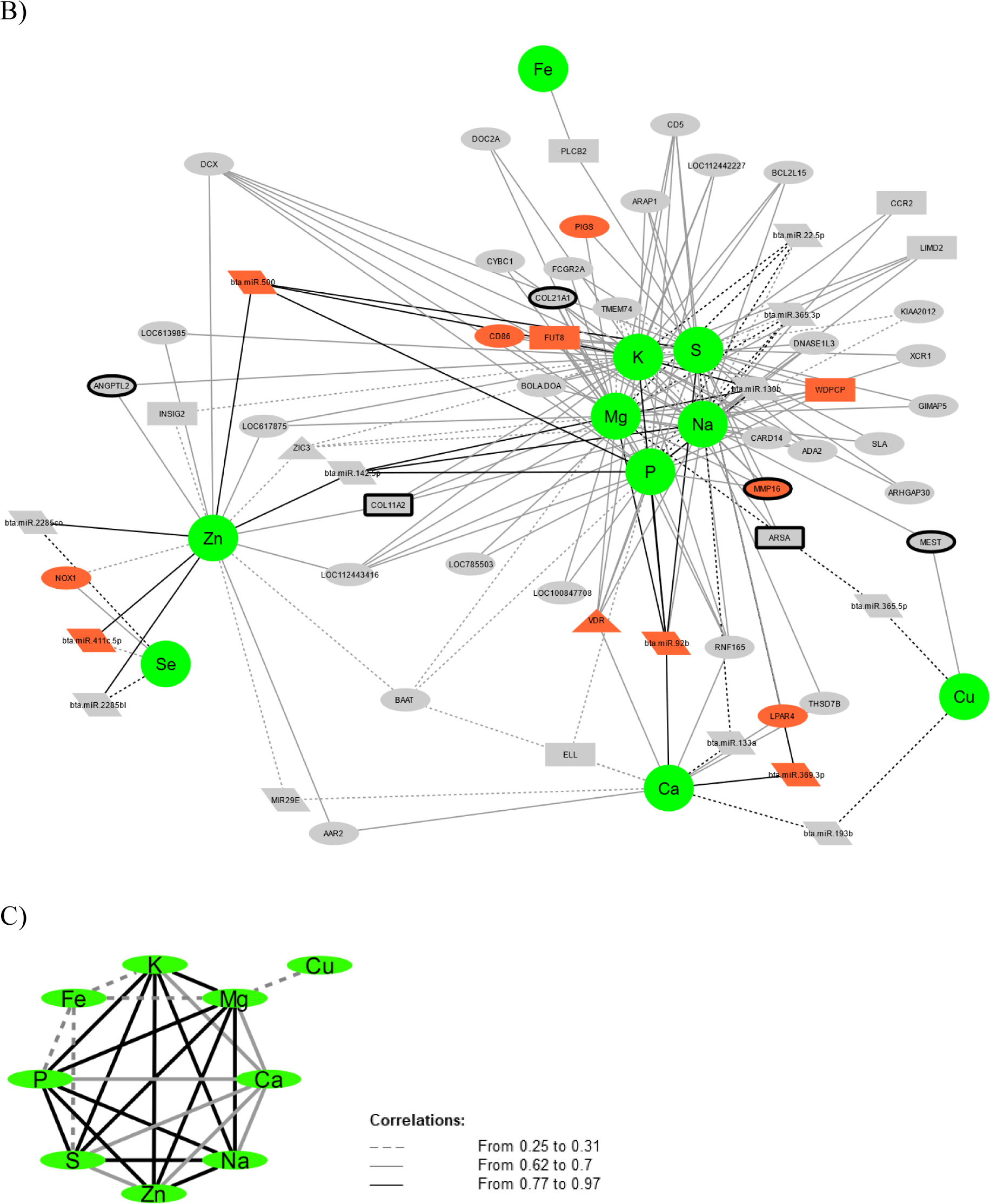
Co-expression network among genes and miRNAs with expression values correlated to at least one mineral. A) Complete network, B) Details about the correlations regarding the genes and miRNAs with expression values correlated to more than one mineral, the internal circle of the complete network, C) Correlations among the mineral’s GEBVs.

**Table 1.** Number of genes and miRNAs with expression values correlated to each mineral considering both PCIT analysis. PCIT general, with mineral genomic estimates of breeding values, genes and miRNAs expression and PCIT miRNA with mineral GEBVs and miRNAs expression. The data came from *Longissimus thoracis* muscle from Nelore steers and the genes and miRNA expressions were identified based on RNA-Seq analysis.

| Mineral | Gene | miRNA | Repeated miRNA<sup>a</sup> |
| --- | --- | --- | --- |
| **Ca** | 22 | 6 | 0 |
| **Cu** | 35 | 5 | 0 |
| **K** | 33 | 5 | 0 |
| **Mg** | 37 | 8 | 0 |
| **Na** | 42 | 6 | 3 |
| **P** | 19 | 6 | 0 |
| **S** | 55 | 6 | 1 |
| **Se** | 32 | 6 | 2 |
| **Zn** | 36 | 9 | 0 |
| **Fe** | 27 | 5 | 1 |

<sup>a</sup> number of miRNAs with expression values correlated to a mineral in both PCIT analysis (PCIT general and PCIT miRNA)

### Principal component score and Regulatory Impact Factor (RIF)

From a principal component analysis based on the GEBVs for each animal, considering ten minerals, we calculated a score for each sample regarding its contribution to phenotypic variation. Based on that, we selected 30 contrasting samples concerning all minerals together, 15 with low score and 15 with high score (Figure 2). These contrasting groups were used to estimate the RIF of all genes and miRNAs with expression values correlated to at least one mineral in the amount of all minerals together, using our application of the original RIF algorithm (see methods). Also, we estimated the RIF of the genes and miRNAs with expression values correlated to each mineral separately using contrasting sample groups for specific minerals. For that, based on the GEBVs, we expanded to 15 the number of samples on the same contrasting groups detailed in previous works with differentially expressed genes regarding mineral amount [9] [10] containing six samples for Ca, Cu, K, Mg, Na, P, S, Se and Zn and five samples for Fe in each group.

**Figure 2.**
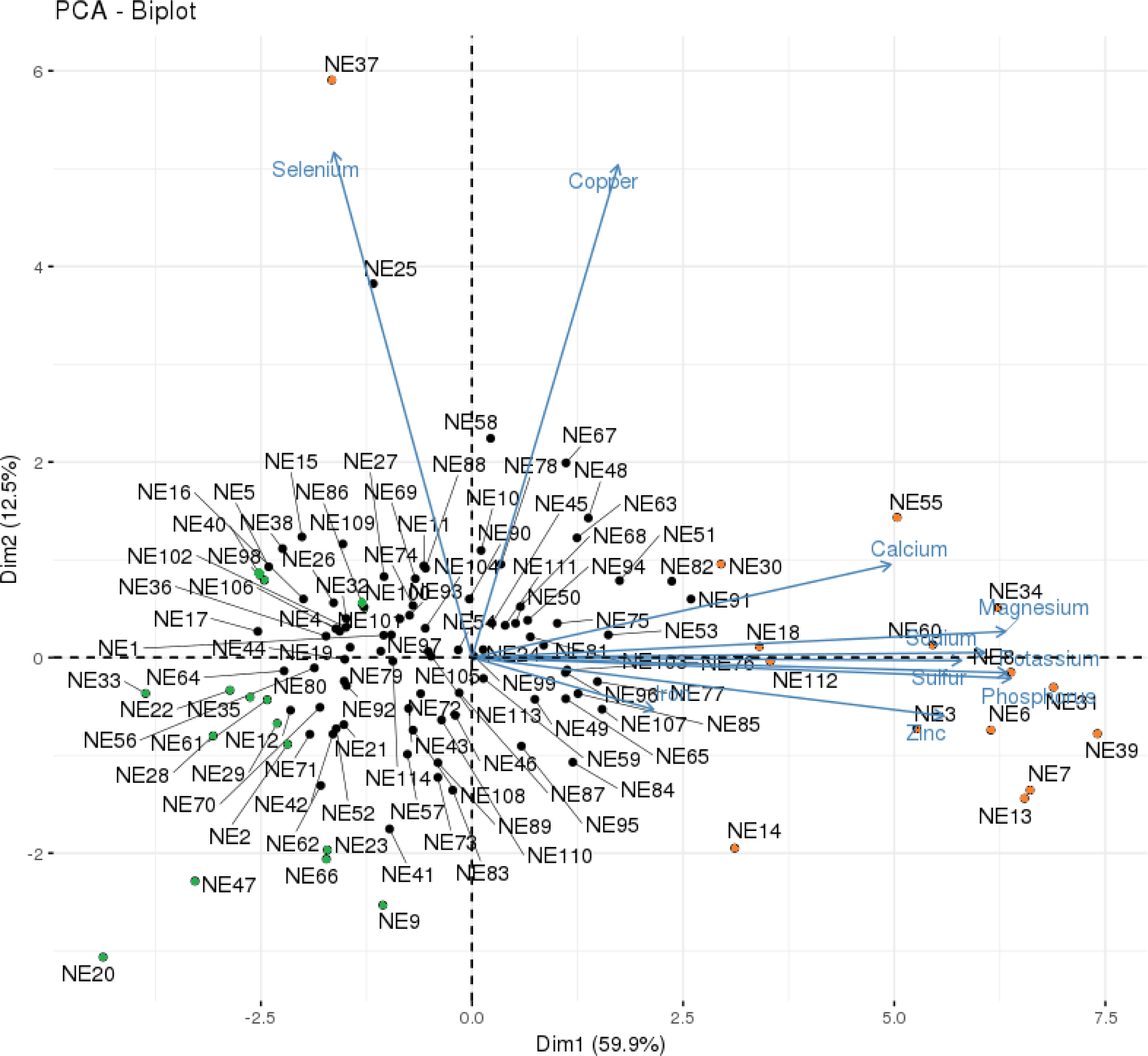
Representation of the contrasting samples considering the genomic estimated breeding values of all 10 minerals together, based on the PCA score. Orange circles represent the samples with the highest scores (positive contrast) and the green circles represent the samples with the lowest scores (negative contrast).

There were 22 genes and two miRNAs with significant RIF based on the high and low score approach. Based on the single mineral analysis, there were three common genes and one common miRNA with significant RIF for two minerals, CD86 molecule (*CD86)* for K and Mg, *VDR* for Mg and Na, WD repeat-containing planar cell polarity effector (*WDPCP)* for Na and P and bta-miR-369.3p for Ca and S. The number of genes with significant RIFs for each mineral varied from zero to seven and for miRNA from zero to two (Table 2). The RIF values of each gene and miRNA presenting significant RIF for each mineral and score analysis is in Supplementary Table S1.

**Table 2.** Number of genes and miRNAs with a significant regulatory impact factor over the genomic estimates of breeding values for each mineral and all minerals together (PCA score). The data came from *Longissimus thoracis* muscle from Nelore steers and the genes and miRNA expressions were identified based on RNA-Seq analysis.

| Mineral | Gene | miRNA |
| --- | --- | --- |
| **Ca** | 1 | 1 |
| **Cu** | 4 | 0 |
| **K** | 3 | 1 |
| **Mg** | 3 | 1 |
| **Na** | 6 | 1 |
| **P** | 1 | 0 |
| **S** | 5 | 2 |
| **Se** | 7 | 0 |
| **Zn** | 4 | 2 |
| **Fe** | 0 | 2 |
| **PCA Score** | 22 | 2 |

### Correlation network

We used the significant correlations between a gene or a miRNA expression and a given mineral, identified in both analyses implemented with the PCIT algorithm, as above described, to derive a co-expression correlation network. To identify potential regulatory mechanisms related to each mineral, we added on this network other layers of information from the same samples, tissue and population, as follows: differentially expressed genes (DEGs) for contrasting mineral amount sample groups [9] [10], transcription factors (TF) [17] and genes affected by eQTLs [18]. This information and genes with significant RIFs were used as node attributes and included in the network analyses (Figure 1). All correlations and attributes necessary to compose Figure 1 are provided (see Supplementary Table S2).

There was at least one putative regulatory element (*i.e.* a significant RIF, TF, miRNA, or gene affected by eQTLs) correlated to each mineral. The number of genes and miRNAs with expression values correlated per mineral per attribute identified is showed in Table 3 and the genes, miRNAs and their attributes are showed in Supplementary Table S2.

**Table 3.**
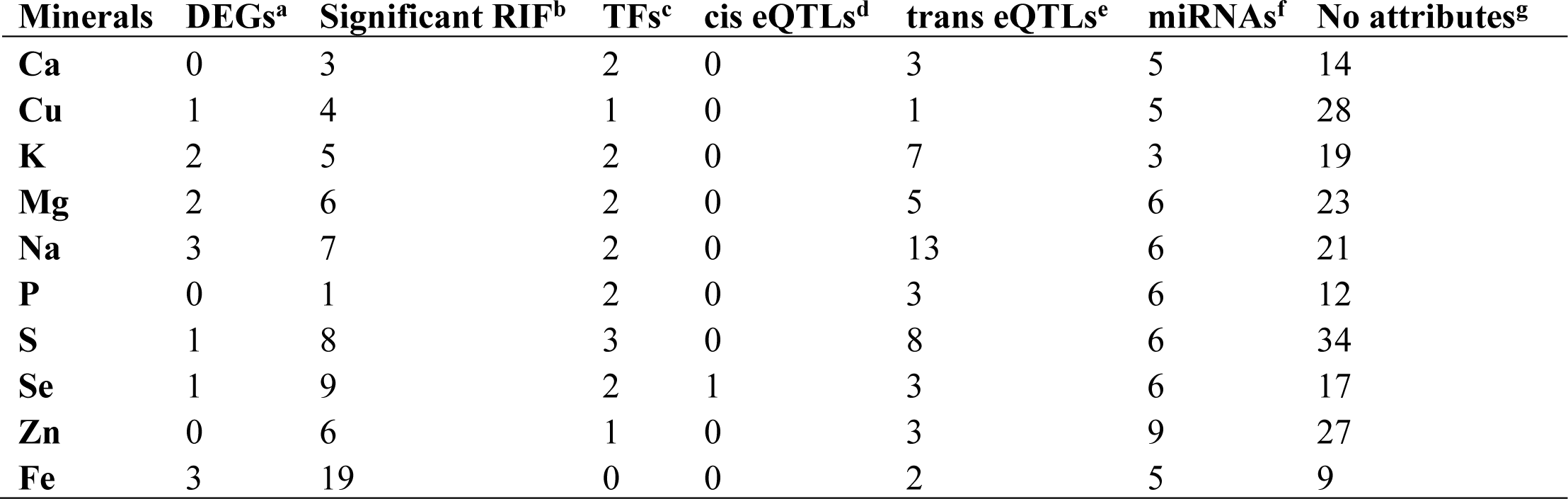
Number of genes and miRNAs with expression values correlated per mineral and per attribute considering both PCIT analysis. PCIT general, with mineral genomic estimates of breeding values, genes and miRNAs expression and PCIT miRNA with mineral GEBVs and miRNAs expression. The data came from *Longissimus thoracis* muscle from Nelore steers and the genes and miRNA expressions were identified based on RNA-Seq analysis. Attributes: a) differentially expressed genes [9] [10], b) genes and miRNAs with significant regulatory impact factor, c) transcription factors [17], d) genes affected by cis eQTLs [18], e) genes affected by trans eQTLs [18], f) miRNAs and g) genes and miRNAs with expression values correlated to each mineral that were not identified in previous works.

There were no functional clusters or over-represented pathways identified in the functional annotation analysis carried out separately for each group of genes correlated to a specific mineral. However, from the functional annotation table, we noted that the gene expressions correlated to the minerals are well conserved among a broad range of organisms. They have functions related to the extracellular matrix, integral membrane constituents, metal ion binding, and partake on regulatory processes linked to transcription, replication, splicing, apoptotic processes, metabolism, transport vesicles, RNA processing, signaling, cell division, adhesion, migration and proliferation, embryonic development and tissue regeneration.

### Integration with differentially expressed genes (DEGs)

To convey the relationship among all genetic elements related to mineral mass fraction detected in our population, we used PCIT to estimate the correlations between a gene or miRNA expression that was found to be correlated to a given mineral in the present work and DEGs previously identified for the same mineral [9] [10]. This analysis was carried out for each mineral separately and included the same genes with regulatory potential as in the previous section (DEGs [9] [10], TFs [17], genes affected by eQTLs [18] and genes with significant RIF). To identify elements with regulatory potential, we then selected the genes that were network hubs or that were significant according to RIF (see methods). We performed a functional annotation analysis with the selected genes for each mineral, separately, to determine which ones were underlying biological pathways.

The expression of all selected putative regulatory elements (hub, significant RIF or miRNA), the ones underlying biological pathways newly identified and the ones being part of enriched pathways in previous work with DEGs related to mineral amount [9] [10] were used as inputs for a final PCIT analyses. This PCIT was carried to identify possible regulators of genes in enriched pathways. Figure 3 shows the co-expression networks built with significant correlations from the final PCIT analyses for Ca, Cu, K, Mg, Na, P, S, Se, and Fe. Supplementary Tables S3 has the correlations and attributes related to creating Figure 3.

**Figure 3.**
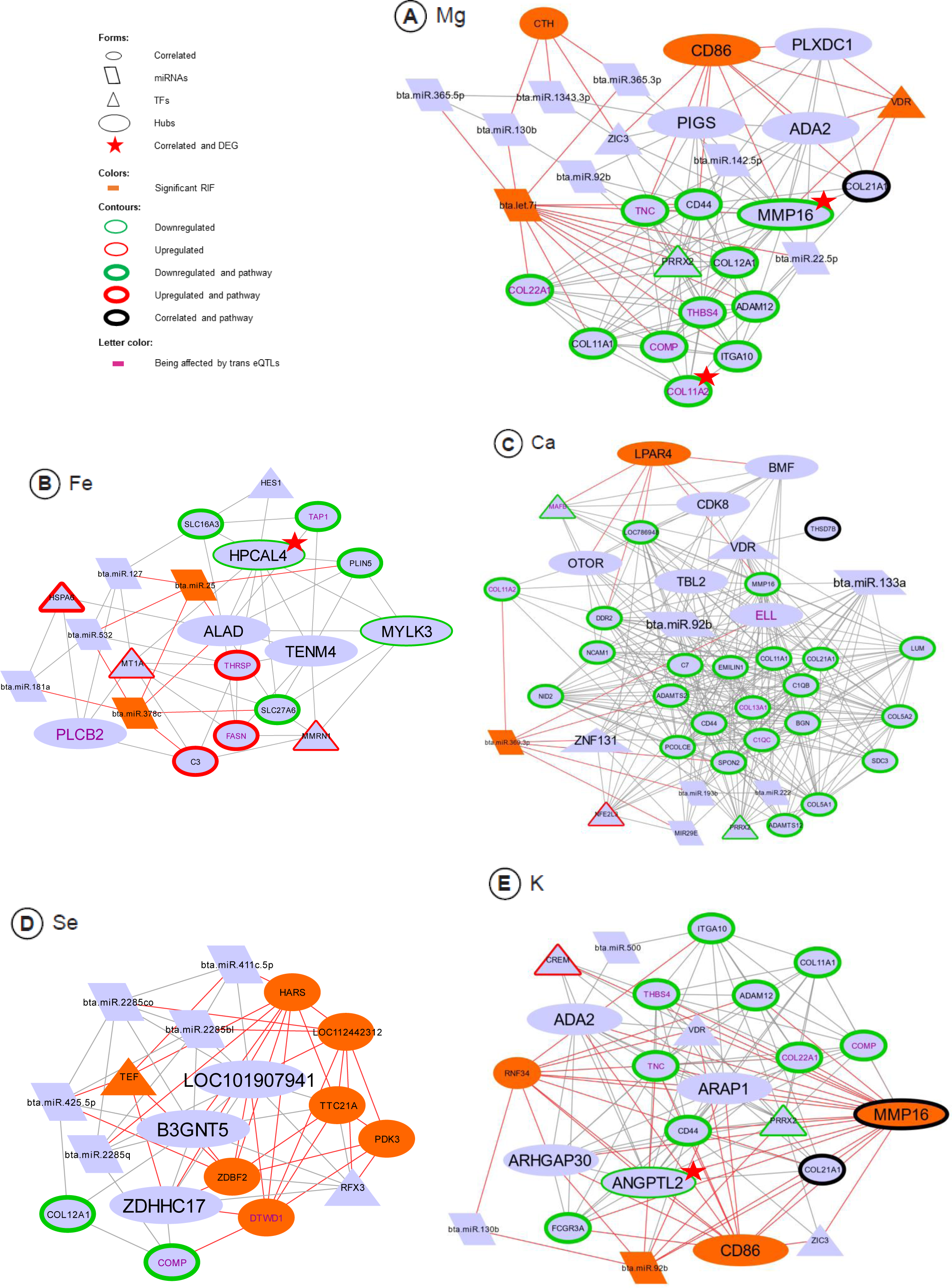
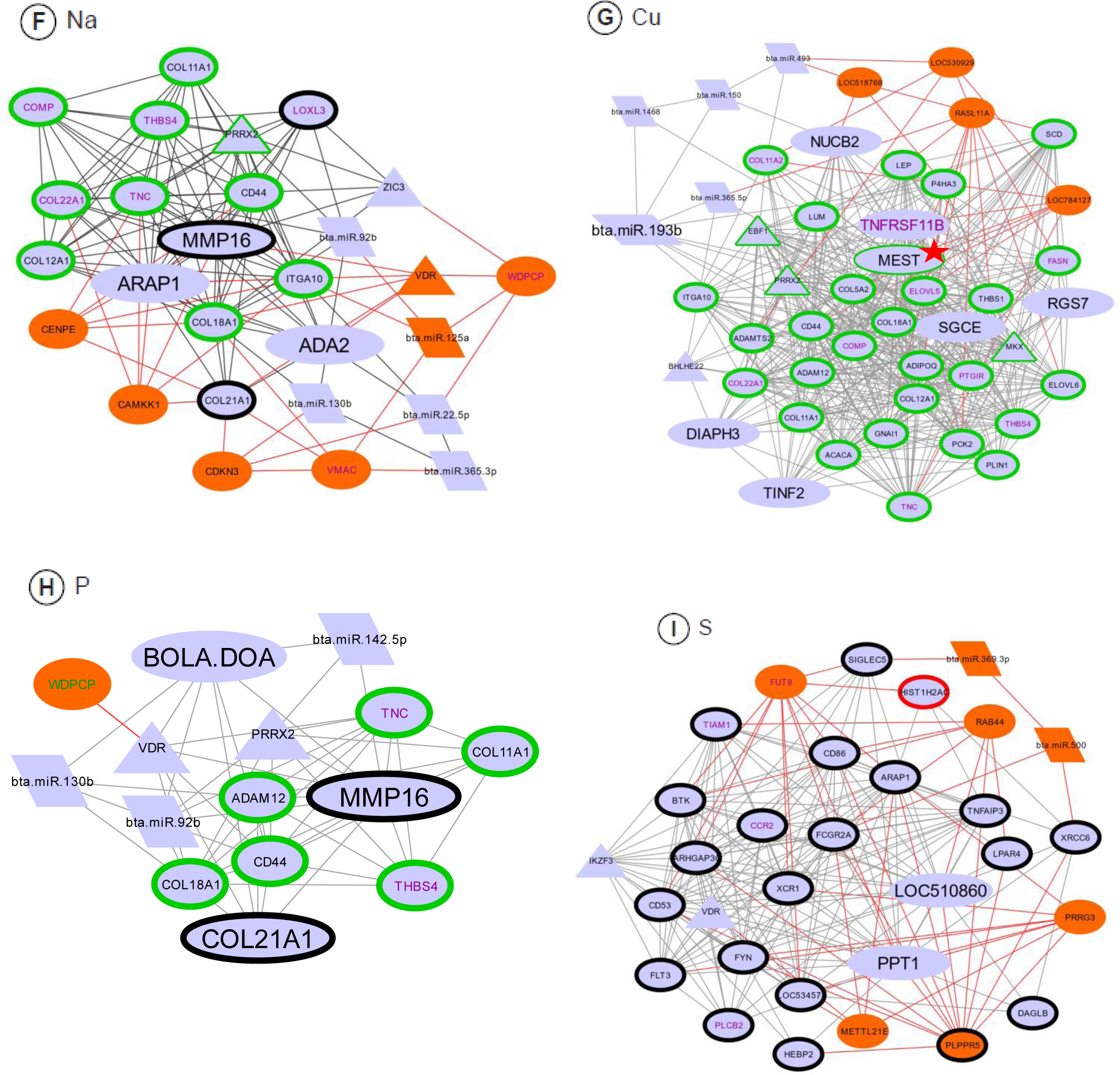
Co-expression networks among genes and miRNAs being part of enriched pathways (DEGs and correlated to a mineral), hubs, TFs, miRNAs or presenting a significant RIF regarding nine of the minerals in study. A) Mg, B) Fe, C) Ca, D) Se, E) K, F) Na, G) Cu, H) P, I) S. Red lines represent the correlations with a significant RIF gene or miRNA.

As we included the differentially expressed genes regarding mineral amount previously detected in in the same population [9] [10], most of the over-represented pathways identified correspond to the previously detected pathways expression analyses. In addition, by the inclusion of correlated genes and pathways from the Reactome database [19], we identified new pathways for K, related to protein metabolism, for Ca, Cu, S and Fe related to immune response, and for S related to signaling. All the pathways enriched for S are new, when compared with our previous work [9]. A list of the pathways enriched for each mineral considering the ones detected with the inclusion of correlated genes expressions and the ones from the previous work [9] [10] is shown in Table 4.

**Table 4.**
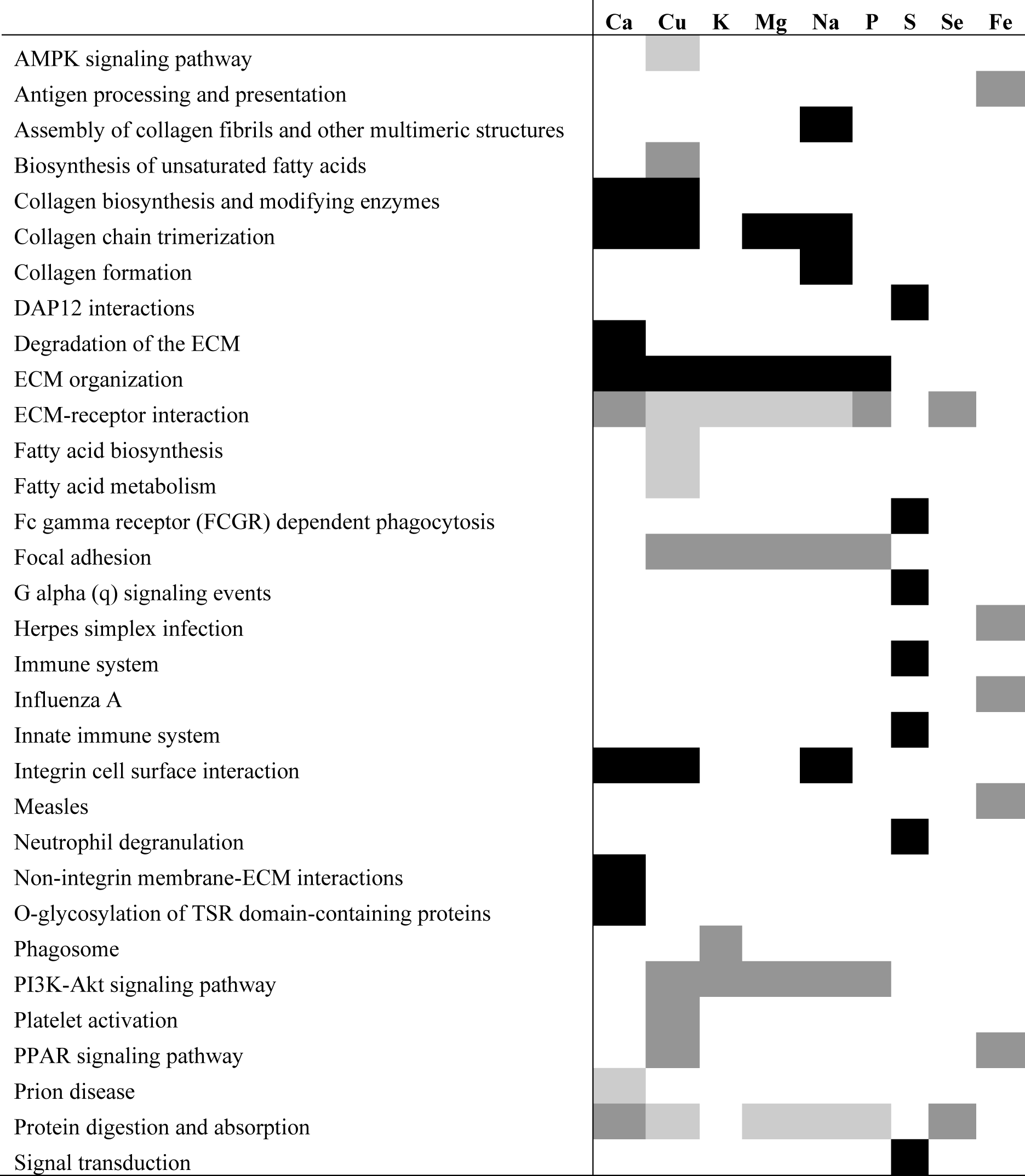
Pathways enriched for each mineral considering the gene expressions correlated to each one of them and the previously detected differentially expressed genes related to the same minerals in the same Nelore population. Pathways just enriched in previous works with a differential expression approach and the same Nelore population are marked in dark grey, pathways enriched in our correlated genes expressions are marked in black and the pathways enriched in both in previous work and in the correlated genes expressions are marked in light grey. There were no enriched pathways for Zn.

Regarding Zn, no gene taking part in the unique enriched pathway previously detected [9] met our criteria. Because of that, for this mineral, we generated a co-expression network by including the DEGs for Zn [9] that had their expression values significantly correlated to hub or RIF elements for Zn and their attributes, in order to identify possible regulators for the DEGs in general. This co-expression network is shown in Figure 4, and the correlations and attributes supporting Figure 4 are presented in Supplementary Table S4.

**Figure 4.**
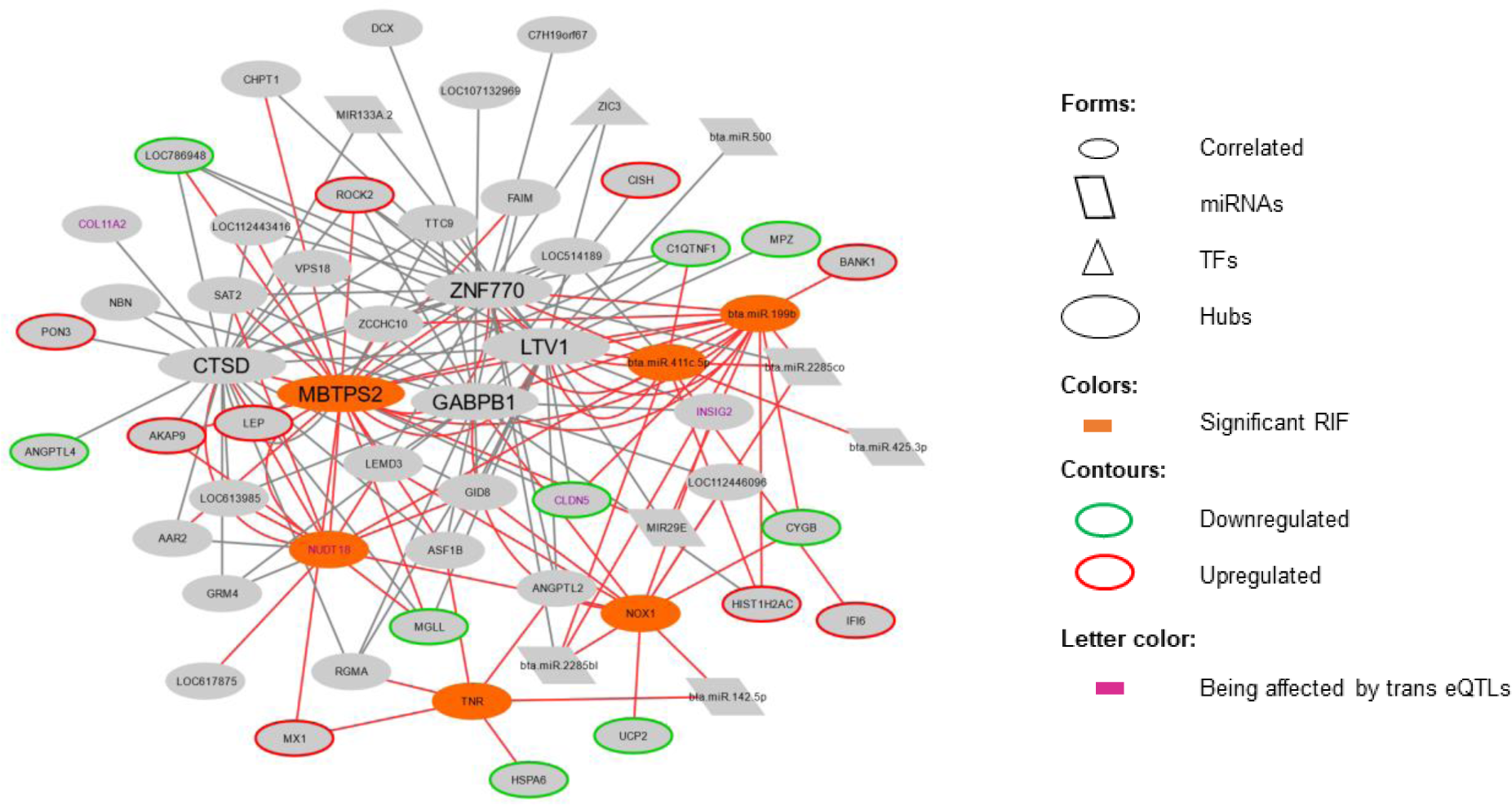
Co-expression network containing DEGs for Zn, genes or miRNAs with expression values that are correlated to these DEGs and are also a hub or a significant RIF for Zn, ora miRNA correlated to Zn. Their functional attributes are presented in different colors or shapes. Red lines represent the correlations with a significant RIF gene or miRNA.

## Discussion

### Relationship among minerals

Correlations identified among GEBVs for most minerals were high (0.77 to 0.97). Thus, a word of caution must inform this discussion of all genes and miRNAs with expression values correlated to each mineral, as correlated responses across minerals may underlie the identified genes and miRNAs, as well as their predicted relationships. All minerals, except Se, were correlated among themselves and all of them revealed genes in common, in the correlation network. In this network, the link between Se and the other minerals was Zn, through the common correlation with the NADPH oxidase 1 (*NOX1)* gene expression, which had significant RIF results for Zn. *NOX1* expression was positively correlated to Zn and negatively to Se. Accordingly, Zn positively regulates NOX1 protein expression in humans, since an increase in Zn leads to a Zn accumulation in the mitochondria. This accumulation increases the production of reactive oxygen species which activates NF-Kb, a known positive transcriptional regulator of *NOX1*, thus increasing its expression [20]. Moreover, Se deficiency is known to induce the oxidation of NrX, a transmembrane protein, by the accumulation of H2O2, which is catalyzed by the NOX1 protein [21]. As the Se deficiency and the H2O2 accumulation catalyzed by the NOX1 protein act in the same known biochemical process, this could explain the negativecorrelation found in our analysis. Further, the oxidation of NrX protein leads to the activation of the Wnt signaling pathway [21], that can act in adult muscle regeneration [22], an evidence for the relevance of this regulation for muscle homeostasis. Another link between Se and Zn were the correlations with three miRNAs expressions: bta-miR-411c-5p (with significant RIF for Zn), bta-miR-2285co and bta-miR-2285bl, although no literature relates these miRNAs to Se or Zn amount, nor to the genes related to these minerals in our analysis.

Fe exhibited a weak correlation with Mg, K, P, and S (from 0.25 to 0.31, p < 0.05) and was linked to other minerals through S, sharing negative correlations with the expression of 1-phosphatidylinositol 4,5-bisphosphate phosphodiesterase gene (*PLCB2)*. PLCB2 protein is critical to Ca efflux [23], although no correlation with Ca amount was found in our data, nor in our previously reported DEGs [9]. The relationship of *PLCB2* gene expression with Fe and S is undocumented, although Fe was reported to cleave the PLCB2 protein in the cornea of bovine, porcine and humans [24]. The *PLCB2* gene is affected by 61 trans eQTLs, harbored across 12 chromossomes [18], making these eQTL regions candidates to regulate this gene expression and consequently Fe and S mass fractions in the muscle.

### PCA score analyses identified regulators of mineral composition

Our score successfully detected contrasting samples regarding all minerals together, allowing for the identification of genes and miRNAs with significant overall RIFs. Considering these genes and the functional enrichment analysis, we identified well-conserved functions for 14 out of 22 genes. From these, we can highlight three with functions related to minerals: Delta-aminolaevulinic acid dehydratase (*ALAD)* encodes a metal ion binding protein linked to Zn, Zinc finger CCHC domains-containing protein 7 (*ZCCHC7),* which encodes a chaperone and Zn finger protein, while Myosin light chain kinase 3 (*MYLK3)* is part of the Ca signaling pathway that participates in muscle contraction.

Mutations in the *ALAD* gene were linked to the phenotypic expression of potentially toxic metal by fly ash exposure in cattle born near thermal power plants, being pointed as a candidate for genomic studies related to metal toxicity [25]. Our results indicated that *ALAD* is a candidate linked to minerals in general, including potentially toxic metals.

### Functional analyses and the search of regulatory elements

Functional annotation analyses, performed based on the genes with expression values correlated to each mineral, showed no functional clusters nor enriched pathways for any mineral. However, some of these genes’ expressions were correlated with DEGs partaking in different pathways and are themselves part of these pathways, which lead us to hypothesize that the remaining genes of the pathways may be modulated in less intensity. This agrees with the small QTL effects already observed for mineral amount [26]. The function annotation for each gene separately showed membrane proteins and extracellular matrix (ECM) related proteins as common annotation for many genes. This observation helps to corroborate the hypothesis that ECM interactions are at the regulatory core for the mineral mass fraction [9]. ECM pathways were enriched for co-expressed groups of genes related to mineral mass fraction and meat quality traits in this Nelore population [13].

When components of a specific pathway are known, a guided-gene approach in a co-expression network can help to identify new genes for the same pathway-related-trait [27], and a pre-selection of genes by biological meaning can improve the network interpretation [12]. Our selection based on enriched pathways, TFs, and significant RIF allowed the inference of genes and miRNAs with a regulatory potential in these pathways. We identified high correlations among these selected elements when compared with the correlations among unselected genes/miRNAs and minerals or considering all genes/miRNAs correlated to a mineral and their respective DEGs. These high correlations and the presence of genes related to regulatory processes reinforces that our methodology can be used to drive the search for meaningful regulatory relationships.

### Potential regulators for more than one mineral

Genes with significant RIF and genes with expression values correlated to others that belong to enriched pathways are the potential regulators. These candidate genes may modulate mineral mass fraction by affecting their target genes and pathways. For the minerals presenting enriched pathways, except Zn, the elements with significant RIFs were connected with miRNAs, correlated genes expressions, TFs and genes being affected by trans eQTLs. They were also part of enriched pathways, reinforcing their regulatory role on the phenotypes. The intricate patterns obtained in these network analyses arise from the fact that the same genes are part of different pathways.

As expected, the pathways identified by considering gene expression correlation with mineral GEBVs were often the same already reported in the differential expression study [9]. The pathways with functions related to ECM processes and protein metabolism were enriched for almost all minerals, except Se, Fe, and Zn. These results also corroborate our previous hypothesis that the regulatory core of mineral amount is linked to ECM processes [9]. Pathways related to fatty acid metabolism were enriched for Cu, as reported in that previous study. However, with the inclusion of the genes with expression values correlated to the minerals, pathways linked to immune responses were enriched for Ca, Cu, Fe, and S. The pathways enriched for S, related to signal transduction and immune response, were not detected in the previous cited work, emphasizing that the integrative approach used herein can bring up new evidences of regulatory processes not identified under the differential expression analysis.

We identified putative regulators that might impact more than one mineral. Cluster of differentiation 86 gene (*CD86)* showed a significant RIF and was a hub gene for Mg and K analyses. The gene *CD86* encodes a protein signaling for T cell activation and proliferation [28] and is linked to T cell adhesion after activation [29]. A Mg sensor, ITK, seems to be required for optimal T cell activation [30] and K^+^ channels are involved in T cell activation, after the binding of the CD86 protein in the CD28 receptor [31], putatively explaining the relationship among these two minerals and *CD86*. The PI3k-akt signaling pathway is activated after this protein-receptor binding in an antigen-presenting cell, leading to downregulation of integrins, participants of the pathways enriched for these two minerals [32]. For both minerals, Mg and K, the known roles of *CD86* support the idea that this is a regulator for the enriched pathways.

The Vitamin D receptor (*VDR*), is a TF with significant RIF for Mg and Na. *VDR* expression has a known relationship with Ca metabolism [33], and it was correlated to this mineral, but it was not identified here as a putative regulator for Ca based on the RIF score. Mg is essential to vitamin D activation, once both enzymes involved in this process, 25-hydroxylase and 1α-hydroxylase, are Mg-dependent [34]. *VDR* expression link with Na is not extensively documented. A putative role of this encoded receptor in the increased Ca absorption and/or reduced Ca loss in menopause women containing no f alleles of the *VDR* gene under a Na and protein-rich diet was reported [35]. The relationship between this gene expression and the ECM processes-related pathways enriched for both minerals seems to be the interaction of the VDR receptor with the Runx2 receptor which, in mammals, stabilizes chromatin remodelers by activating genes involved in ECM mineralization [36].

WD repeat-containing planar cell polarity effector (*WDPCP)* is a gene with significant RIF for Na and P and was affected by one trans eQTL in chromosome five [18]. The *WDPCP* gene encodes a protein that inhibits Wnt activity [37], whose pathway acts in adult muscle regeneration [22], and is activated by high P amounts [38]. ECM processes-related pathways were also enriched for these minerals. ECM stiffness increases the expression of several members of the Wnt pathway through integrins and focal adhesion pathways [39], thus relating the *WDPCP* gene expression with the enriched pathways. The link between *WDPCP* expression and Na is not known. In both minerals, Na and P, *WDPCP* expression value is correlated positively (0.19) with the TF *VDR* expression that represses the Wnt pathway [40].

The miRNA bta-miR-369-3p had a significant RIF for Ca and S. The genes with expression values correlated to this miRNA are not known targets to it. This miRNA expression levels increases in skin and serum of humans with psoriasis [41]. A homolog of psoriasin, a common protein in psoriasis patients, was identified in bovines and have the same antimicrobial and immune response activity as the human one [42]. Psoriasis trigger seems to be the activation of the cellular immune system [43], probably explaining why the bta-miR-369-3p expression level was correlated to several genes involved in immune pathways for Ca and S. Further, Ca and vitamin D play important roles in keratinocyte differentiation and regulate proteins involved in psoriasis [44] and S is used as a known treatment and prevention of recurrence for this disease [45]. Our results suggest the genes expressions correlated to bta-miR-369-3p expression as non-described candidate targets of this miRNA, linked to immune response and mineral concentration.

### Potential regulators for a specific mineral concentration

Some putative regulators showed significant RIF for only one mineral. The miRNA bta-let-7i showed significant RIF for Mg and one of the correlated genes, Collagen alpha-1 (XI) chain (*COL11A1)* is a target of this miRNA. The *COL11A1* gene is a DEG, associated to protein digestion and absorption, as well as, to ECM receptor interaction. This gene encodes a collagen protein, the most abundant protein in ECM. *COL11A1* expression is correlated to Mg, which stimulates collagen synthesis [46], and its expression is correlated to other genes expressions being part of the same or related pathways. Cystathionine gamma-lyase (*CTH)* is also a gene with significant RIF just for Mg. This gene expression is correlated to a Zn finger protein of the cerebellum (*ZIC3)*, a TF, which was correlated to the already mentioned *CD86* gene expression, also associated with Mg herein.

We identified two genes with significant RIF specifically for K: Matrix metallopeptidase (*MMP16)* and E3 ubiquitin-protein ligase (*RNF34)*. The gene *MMP16* encodes a protein whose family is involved in the breakdown of ECM, mostly of collagen genes [47], explaining its link to the enriched pathways related to ECM organization and its correlation with Collagen type XXI alpha 1 chain (*COL21A1)*. Both *MMP16* and *RNF34* expressions were correlated to *CD86* expression, for which the link to K was already discussed. *RNF34* encodes a RINF finger protein that negatively regulates the NOD1 pathway, involved in receptors activating immune responses, similar to *CD86*. Bta-miR-92b expression was correlated to seven genes expressions, and one of them, *MMP16*, is a known target for this miRNA regulation, which could explain the relationship of this miRNA with the over-represented pathways.

For Na, we identified six genes with significant RIF: *WDPCP* and *VDR*, linked to the already discussed ECM processes, Vimentin type intermediate filament associated coiled-coil protein (*VMAC),* Cyclin-dependent kinase inhibitor 3 (*CDKN3),* Centromere protein E (*CENPE)*, and Calcium/calmodulin-dependent protein kinase kinase 1 (*CAMKK1)*. VMAC intermediates filament, play an important role in cytoskeletal organization [48]. Cell adhesions, mediated by integrins, link ECM and cytoskeleton [49]. *CDKN3* encodes a cycling-dependent kinase inhibitor that is involved in cell cycle regulation [50], a process where integrins act [51]. The presence of an integrin gene, integrin subunit alpha 10 (*ITGA10)* in the network, as well as actin interactions, could explain the link of these two genes and the ECM-related pathways for Na. Na presents a miRNA with significant RIF, bta-miR-125a, presenting its expression values correlated to two genes with significant RIF, *WDPCP* and *VMAC,* and the integrin gene *ITGA10*. This miRNA targets *VMAC* who is also affected by six trans eQTLs in chromosome six, being candidates to future studies.

The miRNAs bta-miR-25 and bta-miR-378c had significant RIF for Fe. Their expression values were correlated to each other, to other miRNAs expression and, as with other miRNA found in our results, the genes expressions correlated to them were not described as their targets. Both miRNAs expressions were correlated to *ALAD* gene expression, also a hub gene in the Fe network. Fe amount in the extracellular environment positively affects ALAD protein level and activity [52]. The relationship with the immune response pathways enriched for Fe seems to be in the proteasome involvement in these pathways. ALAD protein modulates proteasome activity [53], and proteasome function can shape innate and adaptative immune responses [54].

Lysophosphatidic acid receptor 4 (*LPAR4)* is a hub gene with significant RIF for Ca, already known to positively regulate cytosolic Ca amount involved in phospholipase C-activating G protein-coupled signaling pathway (GO:0051482). Its expression is linked in our network to MAF BZIP transcription factor B (*MAFB)*expression, a TF that interacts with Gcm2 and modulates parathyroid hormone, which in turn regulates Ca mass fraction [55]. These genes expressions were correlated to other six genes expression. Three of them were DEGs for Ca being part of pathways involved in ECM processes, and the other three were hub genes. From these hub genes, Bcl-2-modifying factor (*BMF)* regulates apoptosis after cell detachment from the ECM [56].

We identified the RAS like family 11 member A (*RASL11A)*, which encodes a RAS protein, with significant RIF for Cu. This gene expression was correlated mainly to the expression of genes involved in fatty acid metabolism, a process where Cu is a known enzymatic co-factor [57]. RAS proteins’ posttranslational modifications are affected by fatty acids [58], possibly explaining the link of this gene expression with the fatty acid-related proteins.

For S, we identified Fucosyltransferase 8 (*FUT8),* RAB44 member RAS oncogene family (*RAB44),* Proline-rich and gla domain 3 (*PRRG3),* Protein-lysine methyltransferase METTL21E (*METTL21E),* and Phospholipid phosphatase related 5 (*PLPPR5)* genes with significant RIF, presenting their expression values correlated or being part of immune response and signal transduction pathways. Sulfur amino acids affect inflammatory aspects of the immune system [59]. Although there is no primary connection between *FUT8* and *RAB44* proteins and the immune system, these proteins contribute to tumor progression [60] [61], in which a robust immune response is involved [62]. *PRRG3* encodes a vitamin K-dependent transmembrane protein with a GLA domain, involved in coagulation factors [63], a process that is linked to the innate immune system [64]. Regarding signal transduction pathways, *METTL21E* was linked to signaling pathways in mouse siRNA experiments [65], and *PLPPR5* encodes a protein member of the phosphatidic acid phosphatase family, acting in phospholipase D mediating signaling [66]. The bta-miR-500, who presented a significant RIF for S is a known regulator of the genes whose mRNA levels were correlated to this miRNA in our analysis.

For Se, all enriched pathways were related to ECM interactions and protein digestion and absorption. For this mineral, we identified six annotated genes with significant RIF, Thyrotroph embryonic factor (*TEF)*, Zn finger DBF-type containing 2 (*ZDBF2),* Tetratricopeptide repeat domain 21 (*TTC21A),* Histidyl-tRNA synthetase (*HARS),* DTW domain containing 1 (*DTWD1),* and Pyruvate dehydrogenase kinase 3 (*PDK3)*. *TEF* is a TF and a leucine zipper protein [67], whose family is required for the activation of DDRs receptors, essential to matrix remodeling [68]. *PDK3* encodes an enzyme responsible for the regulation of glucose metabolism, among many other functions, is related to ECM remodeling [69]. We could not find a link among *ZDBF2*, *HASR,* and *DTWD1* genes expression and Se or the enriched pathways. They are candidates for future studies regarding these potential relationships.

Regarding Zn, even without over-represented pathways, it is possible to infer that the six elements presenting significant RIF are putatively regulators of several correlated genes expressions and a few DEGs, as already discussed by *NOX1*. From the six genes with significant RIF, Membrane-bound transcription factor peptidase, site 2 gene (*MBTPS2)* is also a hub gene encoding an intramembrane Zn metalloprotease and *TNR* encodes an ECM glycoprotein. This information can lead to the assumption that ECM processes can also be associated to Zn amount, as they putatively do to most of the other minerals in study [9].

### New application for PCIT and RIF algorithms

The first co-expression network, containing genes and miRNAs expressions correlated to the mass fraction of at least one mineral, is considered to be a correlation network among elements from two different sources: sequencing (mRNA-Seq and miRNA-Seq) and a measure referring to the trait of interest, the minerals’ GEBVs. Originally, outputs from PCIT algorithm forms co-expression networks based on significant correlations between gene and miRNA expression levels. PCIT works in two steps: first, a partial correlation is calculated for every trio of genes/miRNAs based on the expression values of these elements in a specific set of samples, giving us the strength of the linear relationship between every two items, independent of the third one. In the end, PCIT calculates, for each trio of genes expression, the average ratio of partial to direct correlations. This value is set as the information theory threshold for significant associations, not the same for every analysis, specific for each trio [15]. Statistically, both steps can be used to test the correlation and the significance threshold of other genetic elements, if they vary in the population. Thus, there is no statistical impediment of using PCIT in the way proposed here, to detect genes and miRNAs whose expression values variate in our samples in correlation with the minerals’ GEBVs, as proposed here, since they already represent just the additive genetic effect of the traits [26].

The RIF algorithm was developed to calculate the impact of TFs over a selected list of genes through the expression values of genes and TFs across samples, in two contrasting groups for the studied phenotype (in our case, minerals). This impact factor is calculated in two ways (RIF 1 and RIF 2). RIF 1 gives high scores to TFs that are most differentially co-expressed, highly abundant, and with more expression difference between the groups. RIF 2 gives a high score to TFs for which the expression can predict better the abundance of DEGs [16]. Again, there is no impediment in the analytical methodology to use other genetic information, *e.g.,* GEBVs, since it variates in the population. In our application, we used genes and miRNAs with expression values correlated to at least one mineral in the place of TFs, and GEBVs were used instead of selected genes. In this case, RIF 1 gives a high score to the genes or miRNAs that are most differentially co-expressed, highly abundant and with more expression difference between the contrasting groups (mineral specific groups and score-based groups, separately) and RIF 2 to genes and miRNAs for which the expression can predict better the magnitude of the GEBVs. Together, both new applications can be used to predict genes and miRNAs expressions correlated to mineral mass fraction and to pinpoint which ones have a regulatory impact over mineral amount.

## Conclusion

By using a modification of the PCIT/RIF methodology, we were able to predict regulatory elements related to the mineral amount of ten minerals, indicating over-represented pathways linked to the mass fraction of each mineral and putative regulators that are mineral specific. Our analyses corroborate the link between mineral amounts and the ECM processes, including a relationship with Zn not seen in our previous analysis. In our proposed approach, PCIT can be applied to predict the relationship between gene transcripts or miRNAs and phenotypes, in a genome-wide fashion. Similarly, RIF may predict the regulatory impact of mRNAs and miRNAs levels over phenotypes. This new approach can be applied for any phenotype that is of interest for genomic selection and livestock breeding.

## Methods

Figure 5 contains a flowchart of the steps of our methodology.

**Figure 5.**
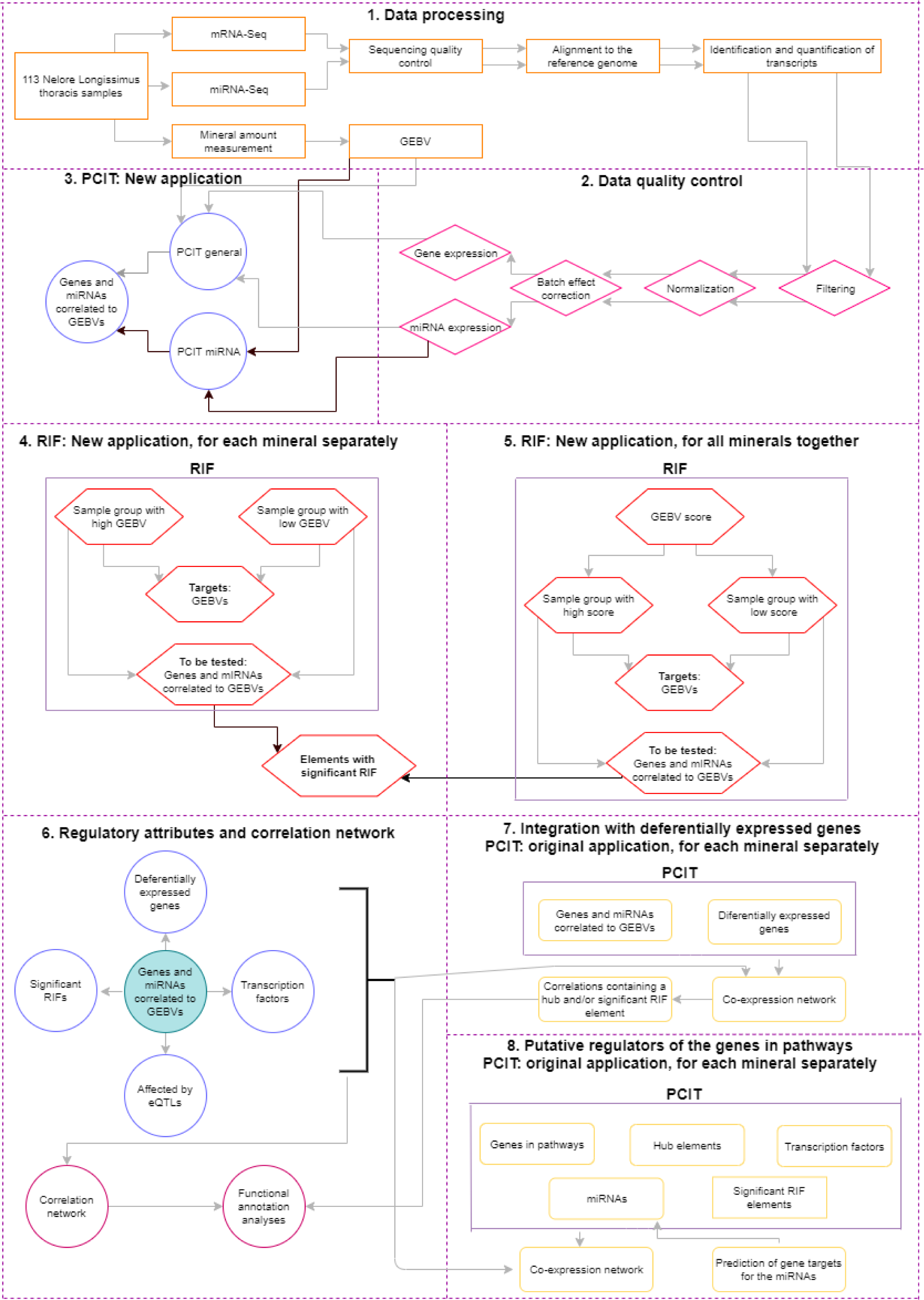
Flowchart representing the steps of the methodology.

### Samples

The Ethical Committee of Embrapa Pecuária Sudeste (São Carlos, São Paulo, Brazil) approved all experimental and animal protocols (CEUA 01/2013). We used the GEBVs from mineral mass fraction [26] and the mRNA-Seq [10], and miRNA-Seq [70] data from 113 samples of *Longissimus thoracis* muscle from Nelore steers that are part of the population already described in previous differential expression analysis related to mineral amount [9] [10].

The animals forming our samples came from a Nelore steer population described elsewhere [26], [71]. In summary, all animals come from half-sibling families, generated by artificial insemination in two different farms, transferred to Embrapa Pecuária Sudeste (São Carlos, São Paulo, Brazil) and maintained in feedlot system with *ad libitum* feed and water access until slaughter, approximately 70 days after the start of the confinement, where the muscle sample collection was done.

### Mineral mass fraction and genetic estimated breeding value (GEBV)

Calcium (Ca), copper (Cu), potassium (K), magnesium (Mg), sodium (Na), phosphorus (P), sulfur (S), selenium (Se), zinc (Zn) and iron (Fe) mass fractions were determined from lyophilized and microwave-assisted digested samples, such as described elsewhere [26]. Calcium, Cu, K, Mg, Na, P, S, Zn, and Fe were determined by inductively coupled plasma optical spectrometry (ICP OES, Vista Pro-CCD with a radial view, Varian, Mulgrave, Australia). Selenium was determined by inductively coupled plasma mass spectrometry (ICP-MS 820-MS, Varian, Mulgrave, Australia).

The estimation of the genetic breeding value (GEBVs) for all the minerals’ amount was previously made [26] through a Bayesian model that considered birthplace, feedlot location and breeding season in the contemporary groups as fixed effects and age at slaughter as a linear covariate.

### mRNA-Seq and miRNA-Seq sequencing and quality control

The total RNA extraction, quality control, and sequencing were described elsewhere [70]. In summary, total RNA from all the 113 samples was extracted using Trizol^®^ (Life Technologies, Carlsbad, CA) and its integrity was evaluated in a Bioanalyzer 2100^®^ (Agilent, Santa Clara, CA, USA). Regarding the mRNA-Seq data, the library preparation was made with the TruSeq^®^ sample preparation kit, and the paired-end sequencing [10] was made in an Illumina HiSeq 2500^®^. For the miRNA-Seq data, the library preparation was made with TruSeq^®^ small RNA sample preparation kit, and the single-end sequencing [70] was made in a MiSeq sequencer.

As a quality control for the sequences, we filtered out reads with less than 65 bp and Phred Score less than 24 for the mRNA-Seq data, and the removal of reads with less than 18 bp and Phred Score less than 28 of the miRNA-Seq data were made using the Seqyclean software (http://sourceforge.net/projects/seqclean/files/).

The reads that passed the quality control were aligned to the reference bovine genome ARS-UCD 1.2 with the STAR v.2.5.4 software [72] for the mRNA-Seq data and with the mirDeep2 software [73] for the miRNA-Seq. The same software was used to the identification and quantification of transcripts and miRNAs, respectively, in raw counts.

### Filtering, normalization and batch effect correction

After quality control, the mRNA-Seq and miRNA-Seq expression data were filtered separately to remove the transcripts and miRNA not expressed in at least 22 samples, or approximately 20% of the samples.

A first component analysis was performed for the mRNA-Seq expression data, with the NOISeq v.2.16.0 software [74] to visually verify the batch effect of the birthplace, feedlot location, breeding season, age at slaughter, slaughter group and a combination of sequencing flowcell and lane over the expression data. The data were normalized using the VST function from DESEq2 software [75], and the batch effect correction for the combination of sequencing flowcell and lane was made using the ARSyNseq function from the NOISeq v.2.16.0 software [74]. For the miRNA-Seq expression data, the procedure was the same, with the batch effect test only for the sequencing lane.

### PCIT (Partial Correlation Coefficient with Information Theory) with mRNA, miRNA and phenotypes

A new application of the PCIT algorithm [15] was developed to test the correlation between the expression values of genes and miRNAs that passed the quality control filters and the GEBVs for ten minerals.

The original application of the algorithm is used to test the co-expression between genes by correlation analysis between expression values [15]. In our application, we included the GEBVs for each one of the ten minerals evaluated here for each sample in the algorithm input with the gene and miRNA expression values (called PCIT general). Using this approach, we estimated the correlations among all the elements. Among the significant correlations, we selected only the genes and miRNAs with expression values correlated to the GEBV of at least one mineral. Due to the low number of miRNAs identified compared to the high number of genes, we did one more PCIT analysis only with miRNAs expression values and the GEBVs (called PCIT miRNA). The results from these two PCITs analysis were combined. In the end we had a list of elements (genes and miRNAs) with expression values correlated to each mineral GEBV.

### RIF (regulatory impact factor)

A new application of the RIF algorithm [16] was applied to obtain the predict regulatory impact of the genes and miRNAs with expression values associated with a given mineral on the amount of the same mineral, considering its GEBVs. The original application of the algorithm was developed to determine the regulatory impact of TFs over selected genes (targets) related to a given trait through their expression values analysis between contrasting groups for the same trait [16]. In our approach, for each mineral, we used the genes and miRNAs with expression values correlated to a mineral, from the previous PCIT analyses, as elements to be tested as regulators and the mineral GEBV as the target.

We carried out 10 different analyses with the RIF algorithm [16], being one for each mineral. As input, we used the GEBVs for the 30 contrasting samples for each mineral as targets (15 representing samples with high mineral mass fraction and 15 with low mineral mass fraction) and the expression values for the genes and miRNAs correlated to the same mineral as elements to be tested. To select these contrasting groups we expanded the sample selection based on GEBVs previously made [9] [10]. Genes and miRNAs with RIF I or II results higher than |1.96| were considered as significant, as authors suggest [16].

### RIF for all minerals together

To identify genes and miRNAs with significant impact factor in all minerals’ mass fraction together, we used the new application for the RIF algorithm [16] using the GEBV from 30 contrasting samples forming two groups regarding the amount of the ten minerals as targets and the expression values for the genes and miRNAs correlated to at least one mineral as elements to be tested.

To select contrasting samples for all the minerals together, we ranked our samples based on a score. To calculate this score for each sample, we performed a principal component analysis (PCA) using the GEBVs for ten minerals for the 113 samples. From the PC results, the score of each sample was calculated based on the following formula:

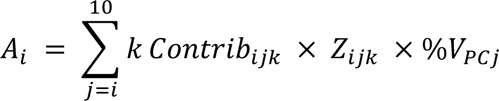

Where: *Ai* = *score* for the animal *i*, 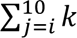 = sum of all minerals k, in all the PCs *j* and in all the animals *i*, *Contrib*_*i,j,k*_ = contribution of the animal *i* in the PC *j* for the mineral *k*, *Z*_*i,j,k*_ = standardized value (standard deviation one and mean zero) of the GEBV for the mineral *k* for the animal *i* in the PC *j* and %*V*_*PCj*_ = eigenvalue of the PC *j*.

We performed a functional annotation analysis using DAVID 6.8 software [76] with the genes presenting significant RIFs for the score, representing all minerals together.

### Genes and miRNAs correlated to minerals

Significant correlations obtained from PCIT [15] analysis between genes or miRNAs expressions and minerals were used to build a co-expression network with the Cytoscape software [77]. We overlapped the gene list from our network with the genes previously reported from our research group based on the same population evaluated here presenting differentially expressed to at least one mineral [9] [10], TFs [17], affected by cis or trans eQTLs [18] and with significant RIF. These features were used as attributes in the network. Regarding the differentially expressed genes (DEGs) for Fe [10], we called the genes more expressed in the high Fe content group as upregulated and the genes more expressed in the low Fe content group as downregulated, to match the nomination of the other minerals’ DEGs [9]. Functional annotation analyses were made using DAVID 6.8 software [76].

### Integration with DEGs

To estimate the relationship among the genes or miRNAs with expression values correlated with minerals and the DEGs between contrasting groups for mineral concentration previously detected [9], we made ten separately PCIT [15] analysis. In these analyses, the PCIT algorithm was used as proposed initially [15] to test the correlations among the genes and miRNAs with expression values correlated to each mineral, and the DEGs previously detected for the same mineral [9] [10].

The significant correlations identified in each analysis was used to obtain co-expression networks with the Cytoscape software [77]. The NetworkAnalyzer tool for the Cytoscape software [77] was used to obtain the connectivity degree of each gene and miRNA in the networks. This value was used to identify the hub genes/miRNAs from the average of the connectivity degree from the network summed with the double of the referent standard deviation.

We considered only the significant correlations containing at least a hub or significant RIF gene/miRNA for a given mineral. The genes present in these considered correlations were used to perform a functional annotation analysis with the STRING v.1.2.2 software [78]. From these analyses, we selected the genes being part of enriched pathways considering KEGG [79] and Reactome [19] databases with *Bos taurus* reference genome.

### Putative regulators of the genes being part of enriched pathways

To identify the elements putatively regulating the genes being part of over-represented pathways for each mineral in the study, we did another round of PCIT [15] analyses, separately for each mineral. In this case, from each mineral last PCIT analysis, we selected as inputs the genes being part of enriched pathways, also considering the previously enriched pathways from differentially expressed genes related to mineral amount [9] [10], the hub elements, TFs [17], miRNAs and the ones with significant RIFs, with their respective attributes. The PCIT [15] results were used to obtain co-expression networks with Cytoscape [77] software.

### miRNA-gene targeting confirmation

We used TargetScan software [80] to predict the target genes for the miRNAs with expression values correlated to a mineral in Figures 3 and 4 and we compared these putative targets with the genes with expression values correlated to them in our networks.

## Acknowledgements

We thank FAPESP (2012/23638-8) for financing the projects encompassing this one and Capes for the scholarship for the first author. We thank all the Staff of Embrapa Pecuária Sudeste responsible for monitoring and taking care of animals. We thank CNPq for the productivity scholarship for the ninth, tenth and last authors. We also thank The University of Queensland for receiving the first author in a Ph.D. internship and the Commonwealth Scientific and Industrial Research Organisation (CSIRO) for assistance during the same internship.

## Authors Contribution

J.A., M.R.S.F., A.R and L.C.A.R. designed the experiments and analysis. J.A., M.R.S.F., A.R., W.J.S.D., A.S.M.C, A.O.L., J.P., M.M.S., L.L.C., G.B.M., A.Z., C.F.G., A.R.A.N., performed the experiments and analysis. J.A., M.R.S.F., A.R. and L.C.A.R. interpreted the results. J.A. and M.R.S.F. drafted the manuscript. All authors revised and approved the final manuscript.

## Competing Interests

The authors claim no competing interests.

